# CD39 REGULATES P2RX7-MEDIATED LUNG NECROTIC LESIONS IN SEVERE EXPERIMENTAL TUBERCULOSIS

**DOI:** 10.1101/2025.05.28.656614

**Authors:** Gislane Almeida-Santos, Igor Santiago-Carvalho, Fabrício Moreira Almeida, Caio César Barbosa Bomfim, Juan Carlo Santos e Silva, Deborah Giovanna Cantarini, Camila Ramos Silva, Martha Simões Ribeiro, Bruna de Gois Macedo, Paulo Henrique Lisboa Raeder, Joaquim Teixeira-Xavier Junior, José Maria Alvarez, Mario Hiroyuki Hirata, Robson Coutinho-Silva, Simon C. Robson, Eduardo Pinheiro Amaral, Elena Lasunskaia, Maria Regina D’Império Lima

**Affiliations:** Universidade de São Paulo (USP), Instituto de Ciências Biomédicas (ICB), Departamento de Imunologia, São Paulo, Brazil; Laboratório de Biologia do Reconhecer, Universidade Estadual do Norte Fluminense Darcy Ribeiro, Campos dos Goytacazes, Rio de Janeiro, Brazil; Inflammation and Innate Immunity Unit, Laboratory of Clinical Immunology and Microbiology, National Institute of Allergy and Infectious Diseases, National Institutes of Health, Bethesda, Maryland, USA; USP, Faculdade de Ciências Farmacêuticas (FCF), Departamento de Análises Clínicas e Toxicológicas, São Paulo, Brazil; Instituto de Biofísica Carlos Chagas Filho, Universidade Federal do Rio de Janeiro, Rio de Janeiro, Brazil; Harvard Medical School, Boston, Massachusetts, USA; Beth Israel Deaconess Medical Center, Boston, Massachusetts, USA; Instituto de Pesquisas Energéticas e Nucleares (IPEN); Carter Immunology Center, Division of Infectious Diseases and International Health, University of Virginia School of Medicine, Charlottesville, VA, USA; Department of Immunology, Mayo Clinic, Scottsdale, Arizona, USA; Institute of Pharmacology and Structural Biology (IPBS), University of Toulouse, CNRS, Toulouse, France

## Abstract

Tuberculosis induces diverse lesions, such as necrotic pneumonia, contributing to disease progression and transmission. Despite advances in understanding the role of ATP-gated P2RX7 ion channels in developing severe forms of tuberculosis, the regulation of this important signaling pathway remains unclear. Herein, we show that the ectonucleotidase CD39 plays an essential regulatory role in TB progression by preventing lung tissue damage, bacterial dissemination, and excessive inflammatory responses. Mechanistically, through its enzymatic activity on the cellular surface, CD39 protects infected macrophages from undergoing necrotic death mediated by P2RX7 activation. We proposed that by protecting macrophages from P2RX7-mediated cell death and bacterial dissemination, CD39 prevents the development of necrotic lesions. Altogether, these findings uncover a significant role for CD39 as an essential component of the molecular regulation underlying the development of severe tuberculosis.

**GRAPHICAL ABSTRACT:** 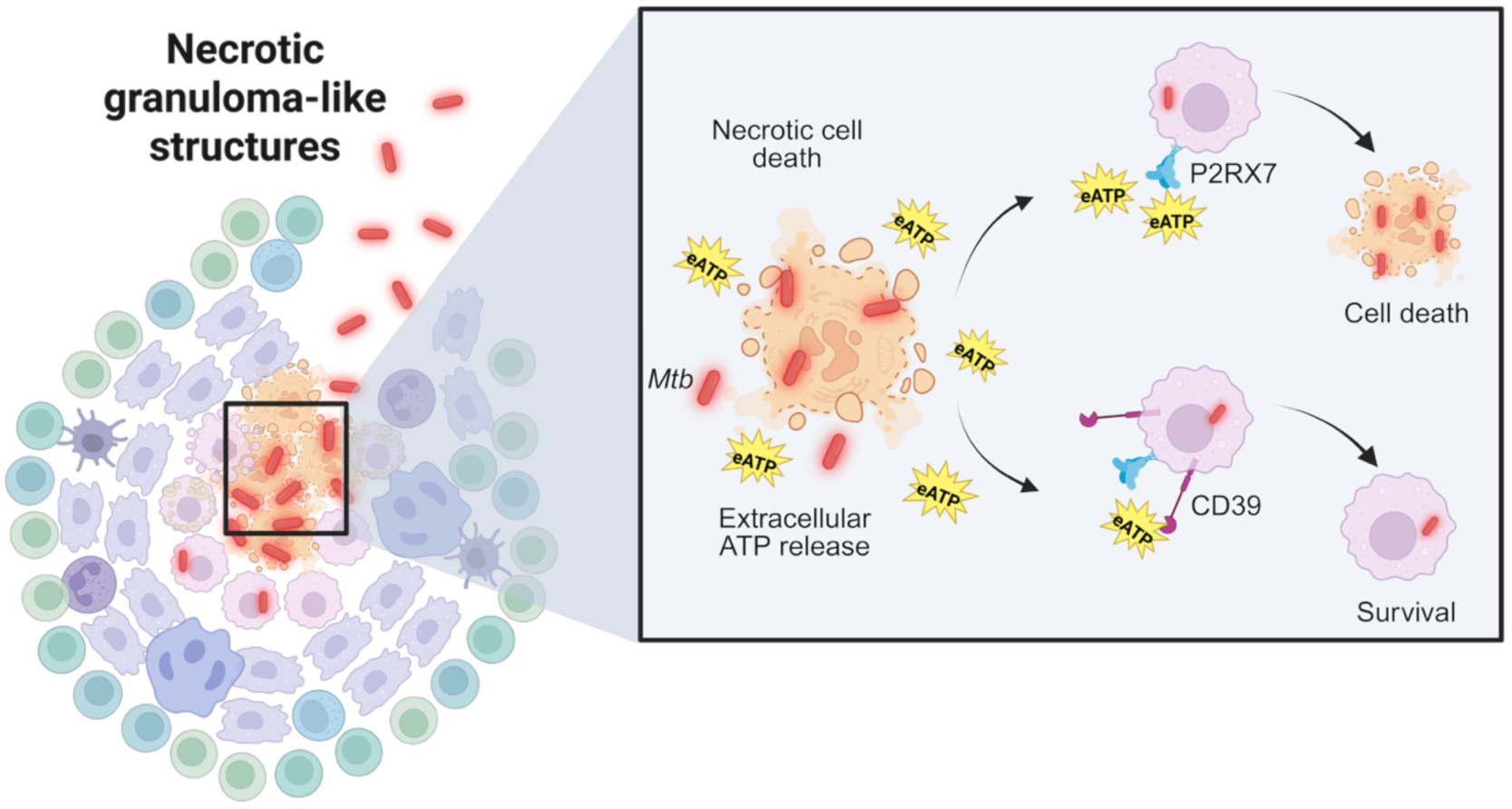

**Brief:** In tuberculosis, necrotic granuloma-like structures release extracellular ATP (eATP), which triggers P2RX7-mediated immune cell death. CD39 degrades eATP, preventing P2RX7 activation and promoting macrophage survival, thereby limiting inflammation, tissue damage, and bacterial dissemination.

## INTRODUCTION

Tuberculosis (TB), a disease caused by *Mycobacterium tuberculosis* (Mtb), continues to be one of the significant global health challenges, with an estimated 10.8 million cases and 1.25 million deaths in 2023 (1). The severe pulmonary form of the TB disease is triggered by the uncontrolled intracellular bacterial multiplication and, consequently, necrotic death of infected myeloid cells, leading to extensive tissue necrosis. Thus, facilitating pathogen transmission (2).

Macrophage necrosis during Mtb infection is considered a pivotal component in promoting disease progression and is detrimental to the host. Unlike apoptosis, where the cell contents are contained, necrosis releases intracellular components, including bacteria, into the extracellular milieu. This process not only promotes the dissemination of Mtb but also exacerbates inflammation by releasing damage-associated molecular patterns (DAMPs), further contributing to tissue damage and the formation of necrotic granulomas (3, 4). Necrotic death in Mtb-infected macrophages has been linked to the activation of multiple regulated necrotic death modalities, such as pyroptosis, necroptosis, and ferroptosis, all of each known to drive exacerbation of the host inflammatory response (5–7). In addition, studies have shown that macrophages undergoing necrosis fail to effectively control bacterial replication, leading to increased bacterial burden in the lungs and worsening tissue damage (8, 9). These combined effects are a hallmark of advanced disease, where the immune system fails to eradicate the infection and instead causes irreversible lung damage (10, 11). Thus, macrophage necrosis in TB represents a detrimental pathway that not only benefits Mtb survival but also drives pathological tissue destruction.

Our previous study has shown that sustained P2RX7 signaling, intensified by high levels of extracellular ATP (eATP), detrimentally affects the progression of TB in mice (12), particularly in cases involving a hyperinflammatory condition induced by hypervirulent mycobacterial strains (13). This evidence suggests that the eATP-P2RX7 axis exacerbates tissue damage under intense cellular stress triggered by the infection. Consistent with this idea, the inhibition of P2RX7 during the acute phase of TB has been shown to improve disease outcome as evidenced by optimal activation of host immune defense against Mtb, non-necrotic granuloma formation, and reduced pathogen dissemination to other organs (14).

Interestingly, the enzymatic activity of CD39 in degrading eATP has been proposed as an essential regulatory mechanism of the immune response during infection and cancer (15–17). In this context, CD39 hydrolyzes ATP to adenosine diphosphate (ADP) and adenosine monophosphate (AMP), which further degrade to adenosine (18). Previous studies have shown that CD39 exhibits a protective effect on preventing liver necrosis by negatively regulating P2RX7-mediated cell death (19). CD39 expression on regulatory T cells and CD4^+^ T cells has been shown to impair macrophage activation and bacterial clearance during TB, contributing to immune evasion (20, 21). Moreover, purinergic signaling through CD39 has been implicated in modulating the balance between pro- and anti-inflammatory responses during *Mtb* infection, which can alter the course of disease progression (22). Although these findings suggest a potential role for CD39 in TB pathogenesis, its direct involvement in regulating macrophage cell death and host immune responses during Mtb infection remains unclear. In this study, we uncovered the effects of CD39 activity on lung immunopathology and its robust association with necrotic lesions during TB. Our data strongly indicate that CD39 expression on immune cells, particularly macrophages, reduces lung necrosis and bacterial dissemination and enhances survival in mice. This underscores the potential of targeting eATP signaling to diminish the widespread necrotic cell death and exacerbated host inflammatory response that worsens TB progression.

## RESULTS

### The expression of genes encoding CD39 and P2RX7 is highly correlated with active forms of TB in mice and human patients

To evaluate the expression of purinergic genes during Mtb infection, we analyzed publicly available datasets of whole and peripheral blood samples of patients with active TB (23–25). Meta-analysis of *ENTPD1* (CD39) gene expression was performed revealing a strong upregulation of the gene compared to healthy subjects in all eight independent reports evaluated (Fig. 1A). Furthermore, enhanced expression of *P2RX7* gene was found in five of these reports, demonstrating a robust correlation between these two genes during Mtb infection. We failed to see differences in the expression of other genes related to the purinergic pathway (ENTPD1-3, NT5E, and ADORA1,2b,3) in patients with active TB. To determine whether this response depends on disease activity, we also analyzed a publicly available RNA-seq dataset (26). *ENTPD1* and *P2RX7* expression was found to be increased in the blood of active TB patients compared to healthy controls and subjects with latent TB infection (LTBI) (Fig. 1B). No significant difference was observed between LTBI and healthy controls, suggesting that *ENTPD1* and *P2RX7* upregulation is associated with active forms of the disease.

**Figure 1.**
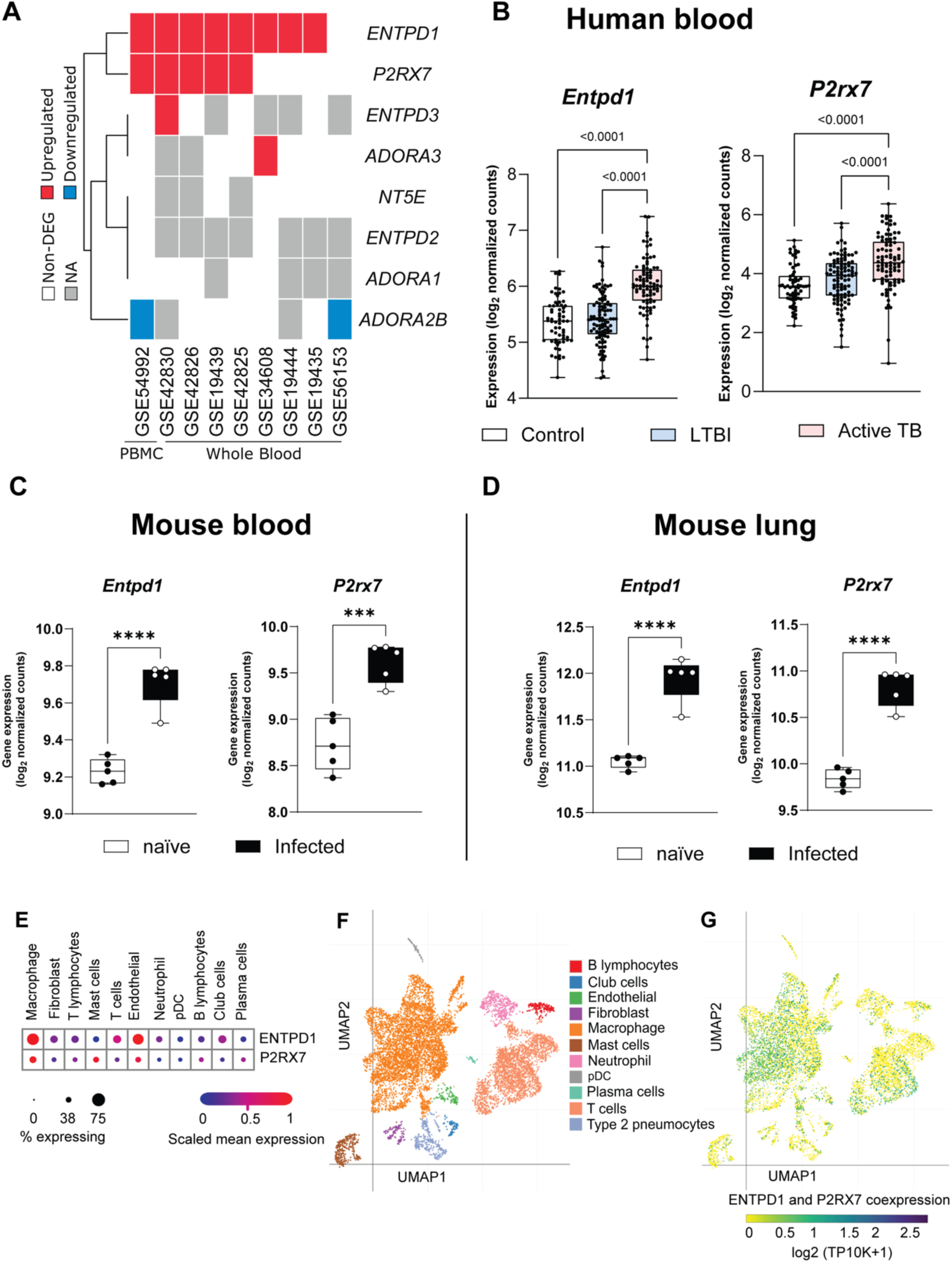
CD39 and P2RX7 gene expression correlate with active tuberculosis (TB) in mice, human patients, and NHP. A) Expression levels of purinergic genes (*ENTPD1*-CD39, *P2RX7*-P2RX7, *ENTPD1-3*, *NT5E, and ADORA1,2b,3*) in whole and peripheral blood samples from patients with active TB were analyzed using publicly available datasets. B) Comparative analysis of *ENTPD1* and *P2RX7* gene expression in blood samples from patients with active TB, latent TB infection (LTBI), and healthy controls derived from publicly available RNA-seq datasets. C & D) Transcriptional levels of *Entpd1* and *P2rx7* (CD) of mice infected with *Mycobacterium tuberculosis* (H37Rv Mtb strain) compared to non-infected (naïve) mice, using publicly available RNA-seq datasets. E) Cell Type Annotation analysis in granulomas from non-human primates, based on public high-throughput single-cell mRNA sequencing datasets. F & G) UMAP plots showing the distribution of immune cell populations (F) and the co-expression of *P2rx7* and *Entpd1* across these populations (G) in granulomas from non-human primates.

To determine whether the transcriptional signature observed in humans is recapitulated in mouse models, the mRNA expression of *Entpd1* and *P2rx7* genes were analyzed in publicly available datasets of the blood and lungs of C57BL/6 (WT) mice infected with mycobacteria (26). The transcriptional mRNA levels of *Entpd1* and *P2rx7* genes were found augmented in both blood and lungs of Mtb-infected mice compared to naïve animals, similarly, to seen in human disease (Fig. 1C and 1D). These findings suggest the upregulation of genes encoding CD39 and P2RX7 as biomarkers for active TB and more specifically indicate their use to infer the development of severe disease.

To overview CD39 and P2RX7 gene expression across different cell subtypes in the infected lung, we analyzed a public high-throughput single-cell mRNA sequencing on granulomas from non-human primates (NHP) at 4 weeks p.i. (27). A Cell Type Annotation analysis showed a higher mRNA expression of *ENTPD1* on endothelial cells and macrophages than on other immune and non-immune cells (Fig. 1E). Interestingly, *P2RX7* expression was slightly increased on endothelial cells, macrophages and mast cells within the granuloma (Fig. 1E). Importantly, among all cell subtypes, macrophages were the most abundant population (Fig. 1F) and the cell type showing the higher *ENTPD1* and *P2RX7* coexpression (Fig. 1G).

### CD39 deficiency aggravates pulmonary necrosis caused by hypervirulent mycobacteria

We previously defined that P2RX7 signaling promotes the development of necrotic lesions in mice infected with highly virulent mycobacteria (12). To investigate the role of CD39 on severe TB, we infected *Entpd1*^−/-^ and their WT littermates with the highly virulent Beijing M299 Mtb clinical isolate due to its ability to trigger necrotic lung pathology in WT mice. (12, 28). *Entpd1*^−/-^ mice had anticipated and accentuated weight loss compared to WT mice (Fig. 2A). Additionally, all *Entpd1*^−/-^ mice died within 18 to 27 days p.i. (Fig. 2B). At the same time, 87.5% of the WT mice remained alive during a 42-day p.i. period. Lungs collected 21 days p.i. showed 200-fold increase in the bacterial burden in *Entpd1*^−/-^ mice compared to WT mice (Fig. 2C). Notably, increased presence of lung white nodules was observed in *Entpd1*^−/-^ mice compared to WT mice (Fig. 2D). Histopathological analysis of the lungs demonstrated extensive necrotic lesions and areas of alveolitis and edema in *Entpd1*^−/-^ mice (Fig. 1E and 1F). In contrast, initial necrotic areas surrounded by alveolitis were observed in WT mice. These results suggest that the expression of CD39 suppresses the development of severe lung disease as evidenced by reduced necrotic pneumonia and mortality in mice infected with highly virulent Mtb.

**Figure 2.**
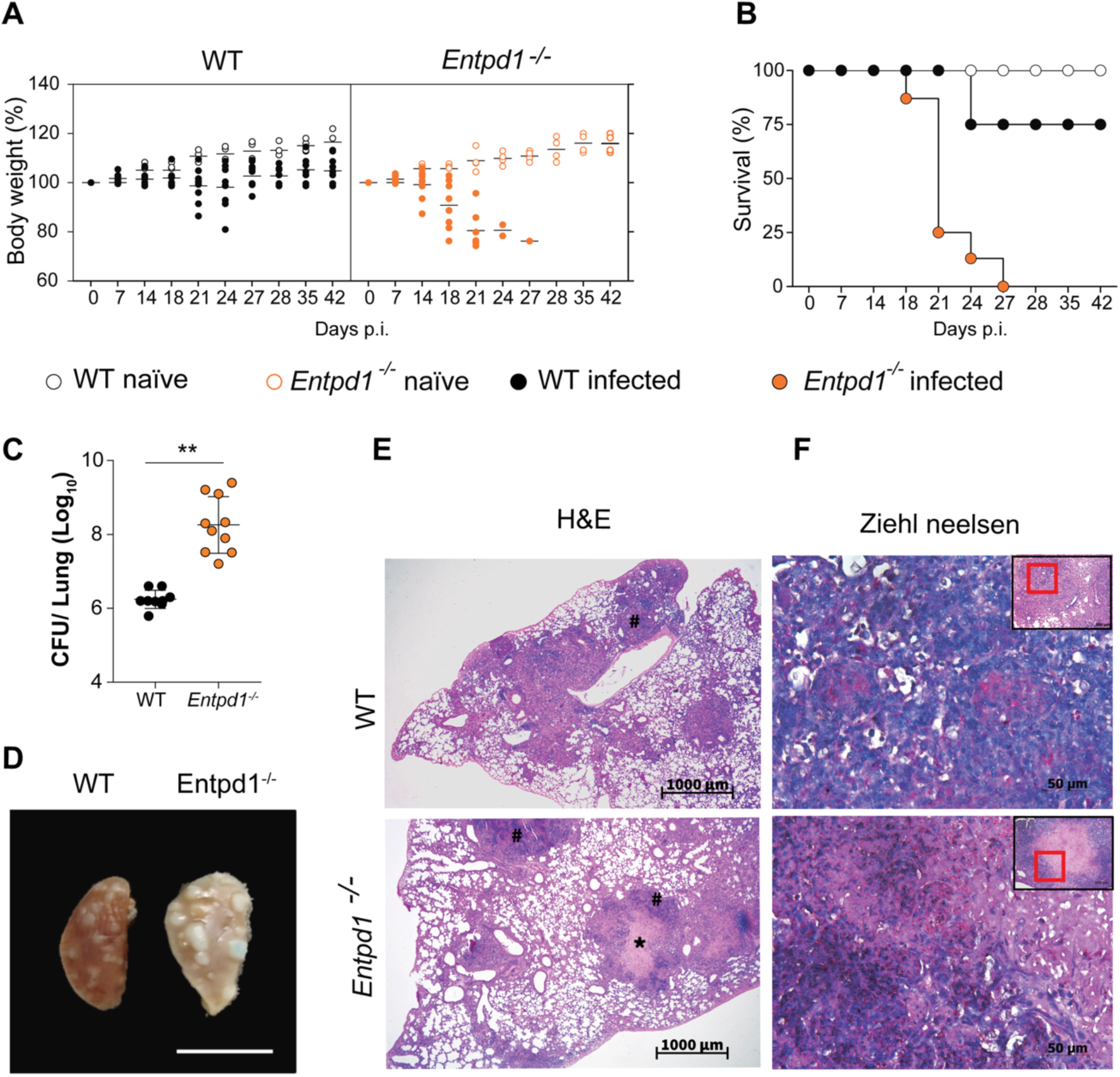
CD39 deficiency exacerbates pulmonary necrosis induced by hypervirulent Mycobacterium tuberculosis (Mtb). C57BL/6 wild-type (WT) and CD39-deficient (Entpd1^−/-^) mice were infected with the hypervirulent M299 Mtb strain. A) Percentage change in body weight relative to baseline (day 0). B) Survival curve of infected mice, where animals were euthanized upon reaching a >20% loss in body weight to comply with humane endpoint criteria. C) Colony-forming units (CFU) in lung homogenates. D) Gross macroscopic pathology of the left lung at day 21 of infection. E) Hematoxylin and Eosin staining of lung sections. F) Ziehl-Neelsen staining of lung sections. Necrotic spots are marked with an asterisk (*), and areas of alveolitis are indicated by a hash symbol (#). Significant differences between groups are indicated by p < 0.05, as determined by the Mann-Whitney non-parametric test. Data represent results from two independent experiments, with five mice per group.

### The expression of CD39 in the immune cells protects against severe TB

To understand if the protective effect of CD39 activity was associated with its expression on immune or epithelial cells, we next evaluated the protein levels of CD39 and P2RX7 on immune (CD45^+^) or non-immune (CD45^−^) lung compartments of WT mice following M299 Mtb infection. The results revealed the accumulation of immune cells in the infected lungs (Fig. 3A), which showed increased proportion and expression of CD39 and P2RX7 compared with non-immune cells (Fig. 3B and 3C). Next, we evaluated whether the expression of CD39 in the immune compartment was required for protection against severe TB using *WT>CD45.1* (WT) and *Entpd1^−/-^>CD45.1* bone marrow mouse chimeras. *Entpd1^−/-^>CD45.1* mice were more susceptible to infection and showed accentuated body weight loss at day 21 p.i. compared to *WT>CD45.1* (Fig. 3D). The lung weight and bacterial burden in *Entpd1^−/-^>CD45.1* mice were also increased compared to *WT>CD45.1* mice (Fig. 3E and 3F). Enlarged lung white nodules were found in infected *Entpd1^−/-^>CD45.1* mice (Fig. 3G), which showed larger necrotic lesions with extracellular bacteria compared to *WT>CD45.1* mice (Fig. 3H). Together, the results suggest that the expression of CD39 on hematopoietic compartment is critical to control the development of necrotic lesions and bacterial spread.

**Figure 3.**
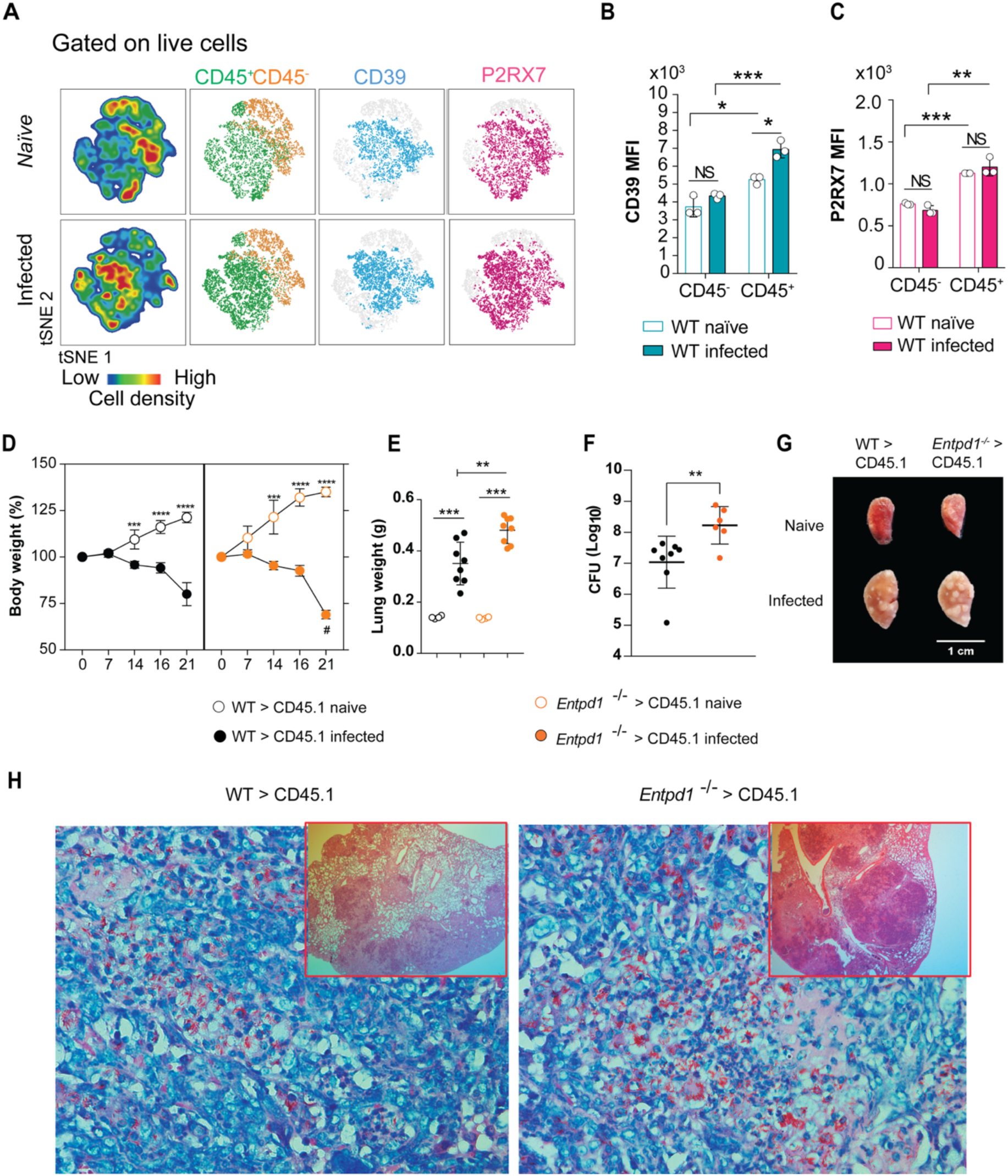
The expression of CD39 in the immune cells protects against severe TB. C57BL/6 (WT) mice, as well as WT>CD45.1 (WT) and *Entpd1^−/^*^−^>CD45.1 bone marrow mouse chimeras, were infected with the M299 Mtb strain. A) Distribution of immune (CD45+) and non-immune (CD45-) cells in the lungs of naïve and infected WT mice. Mice were analyzed on day 21 of infection. B & C) Median intensity fluorescence (MFI) of CD39 and P2RX7 expression in CD45+ and CD45-lung cells described in A. D) Percentage changes in body weight relative to baseline (day 0) in infected mouse chimeras. E & F) Lung weights and colony-forming units (CFUs) in lung homogenates of infected mouse chimeras on day 21 of infection. G) Gross macroscopic pathology of the left lung in mouse chimeras on day 21 of infection. H) Histopathological analysis with Ziehl-Neelsen staining of lung sections of mouse chimeras at 21 days of infection. Significant differences between groups are indicated by p < 0.05, when * represents p < 0.05, ** p < 0.01, *** p < 0.001, as determined by the Mann-Whitney non-parametric test. The data represent results from two independent experiments, with five mice per group.

### The ablation of CD39 results in lower numbers of macrophages and dendritic cells while increasing the inflammatory response in infected lungs

To investigate which immune cell population could be responsible for the protective role of CD39 in preventing TB severity, we evaluated the expression of CD39 and P2RX7 in macrophages, neutrophils, monocytes, dendritic cells and T cells from the lungs of infected WT mice. Macrophages exhibited the highest expression levels of both molecules (Fig. 4A and 4B), as observed in the single-cell analysis of granulomas from non-human primates (Fig. 1E-G).

**Figure 4.**
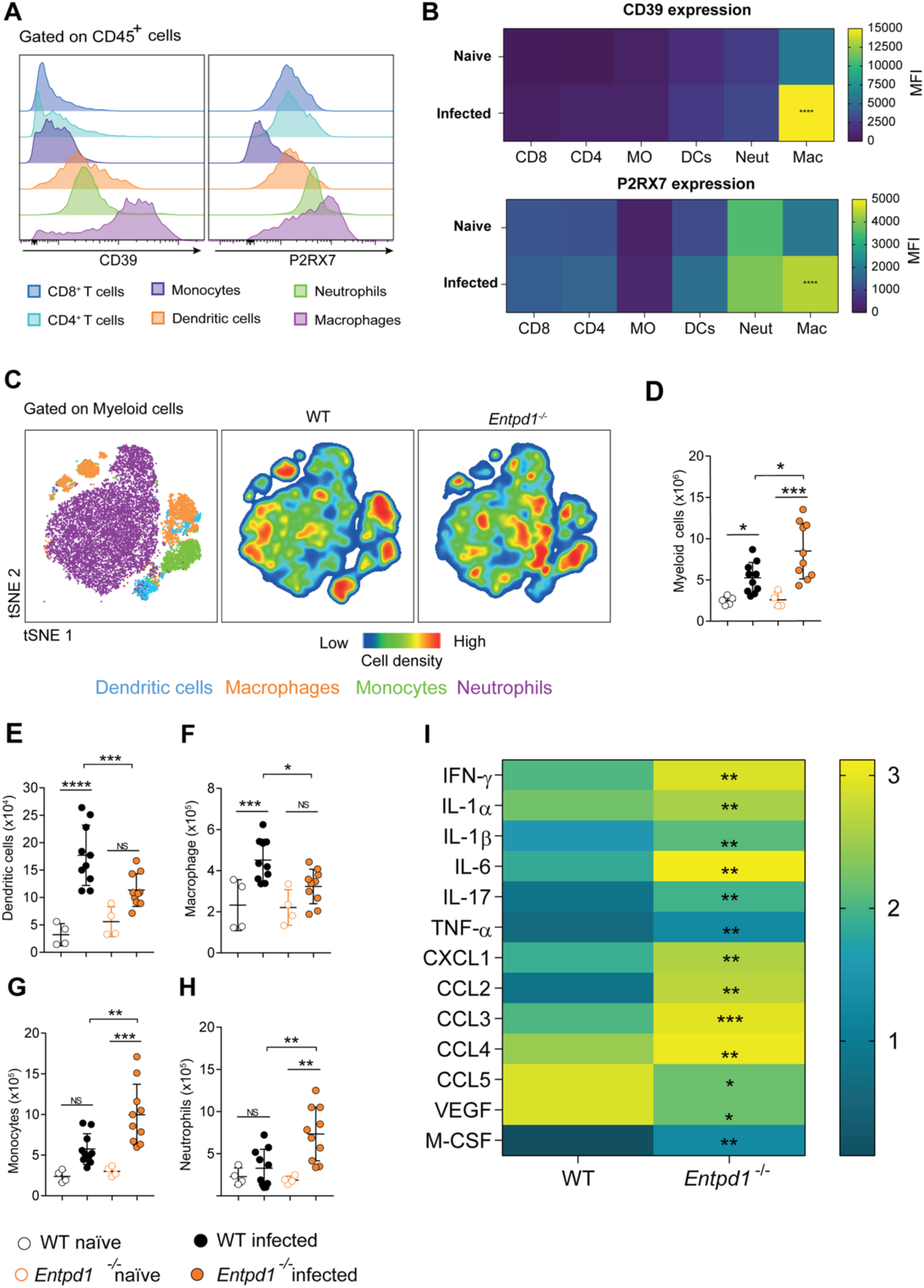
CD39 deficiency reduces macrophage and dendritic cell populations, promoting inflammation in the Mtb-infected lung. A) Histograms showing a qualitative comparison of CD39 and P2RX7 protein expression levels across immune cell subsets isolated from the lungs of Mtb-infected WT mice. B) Heatmap illustrates the Geometric Mean Fluorescence Intensity (GMFI) of P2RX7 and CD39 across immune populations in the infected WT lungs. C) tSNE plots depicting the distribution of macrophages (Live/Dead⁻, DUMP⁻, CD45⁺, Ly6G⁻, CD68⁺, CD64⁺), dendritic cells (Live/Dead⁻, DUMP⁻, CD45⁺, Ly6G⁻, CD68⁻, CD64⁻, CD11b⁺, MHCII⁺), monocytes (Live/Dead⁻, DUMP⁻, CD45⁺, Ly6G⁻, MHCII⁻, SSCLow, Ly6C⁺), and neutrophils (Live/Dead⁻, DUMP⁻, CD45⁺, Ly6G⁺) within the Myeloid cell (Live/Dead⁻, DUMP⁻, CD45⁺) population. D-H) Absolute numbers of myeloid cells, dendritic cells, macrophages, monocytes, and neutrophils in infected and non-infected WT and Entpd1^−/-^ mice. I) Cytokine and chemokine levels in lung homogenates of infected WT and Entpd1^−/-^ mice. Significant differences between groups are indicated by p < 0.05. Significance levels are represented as follows: p < 0.05 (*), p < 0.01 (**), p < 0.001 (***), and p < 0.0001 (****), determined by the Mann-Whitney non-parametric test. The data represent results from two independent experiments with five mice per group.

Next, we investigated the impact of CD39 deficiency on the inflammatory response in WT and *Entpd1*^−/-^ infected lungs. We have previously demonstrated that CD4^+^ T cell response in the lung parenchyma plays an essential role in the severity of TB caused by highly virulent strains (29). We observed decreased numbers of CD4^+^ T cells in the lung of *Entpd1*^−/-^ mice compared to WT mice. However, there were no differences in the numbers of parenchymal CD44^+^CD4^+^ T (CD45 i.v.^−^) or intravascular (CD45 i.v.^+^) CD44^+^CD4^+^ T or CD44^−^CD4^+^ T cells when comparing infected WT and *Entpd1*^−/-^ mice (Sup. Fig. 1). Despite the increase in the absolute CD11b^+^ number per lung of myeloid cells in infected *Entpd1*^−/-^ mice (Fig. 4C) the absolute number per lung of macrophages and dendritic cells were reduced compared to infected WT mice (Fig. 4D-F). At the same time, neutrophil and monocyte populations augmented in the absence of CD39 (Fig. 4D, 4G and 4H). In addition, we observed increased levels of cytokines implicated in the necrotic process (IL-1α and IL-1β), cell migration (CCL2, CCL3, and CCL4), and effector T cell response (IFN-ψ, TNF-α, IL-17) in lung homogenates of infected *Entpd1^−/-^* mice (Fig. 4I). These findings suggest that the lack of CD39 leads to the depletion of macrophages and dendritic cells while promoting the inflammatory response against TB.

### CD39 deficiency leads to an increased macrophage death and inflammasome activation in Mtb-infected *Entpd1^−/-^* mice

To assess whether the low macrophage numbers and high IL-1β levels in the lungs of infected *Entpd1^−/-^* mice were associated with increased cell death and inflammasome activation, we analyzed the lung myeloid cell population according to cell viability and caspase-1 activation. A percentage reduction of live cells (Live and Dead^−^) and increase of active caspase-1^+^ dead cells (Live and Dead^+^) was observed in infected *Entpd1*^−/-^ mice compared to infected WT mice (Fig. 5A-C), while there was no difference in the active caspase-1^+^ live population (Fig. 5A-D). Of note, a population of dead cells expressing high levels of active caspase-1 became evident only in the absence of CD39. These results indicate that CD39 prevents inflammasome activation and myeloid cell death, possibly by a mechanism dependent on pyroptosis.

**Figure 5.**
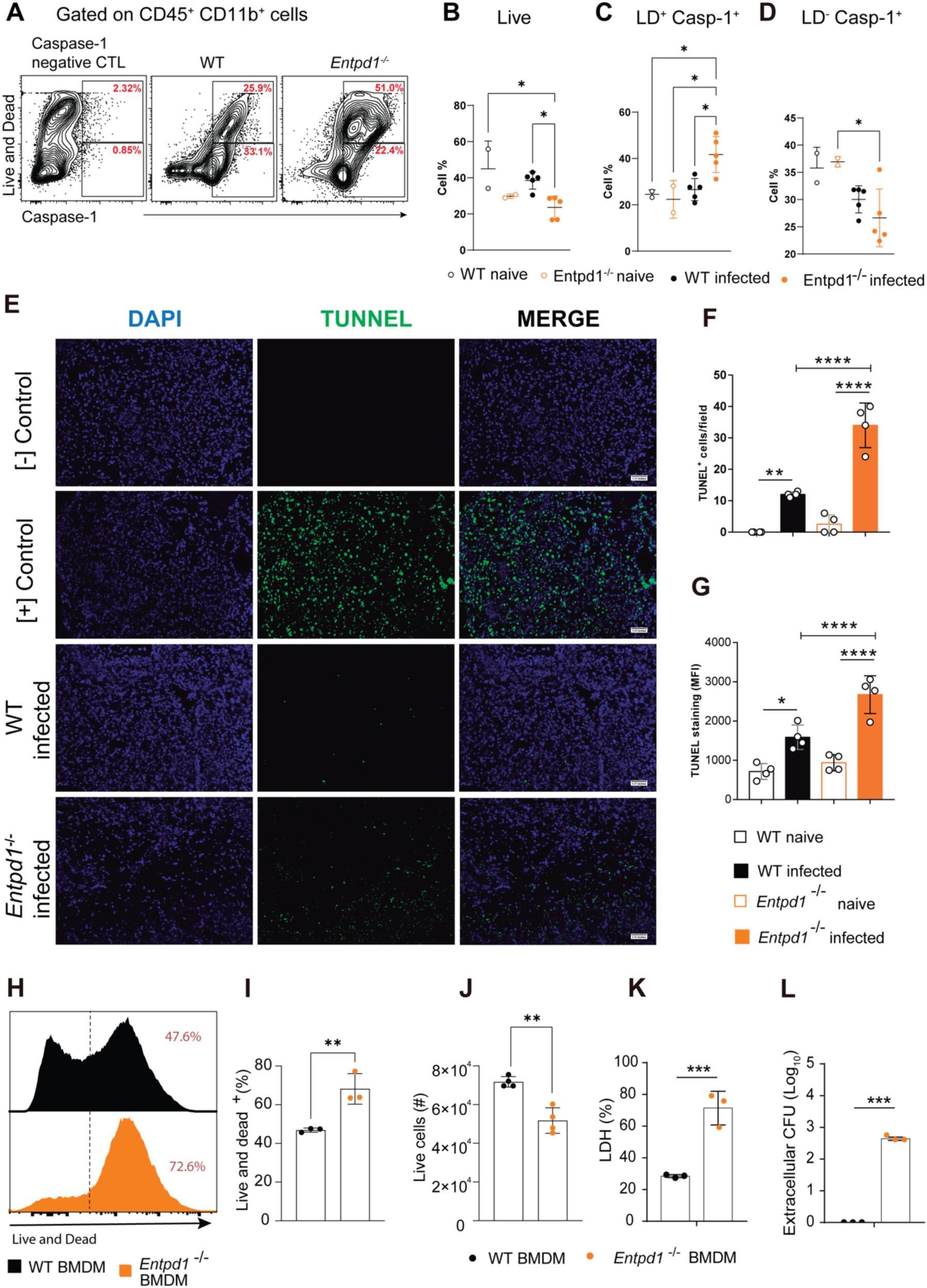
CD39 expression prevents caspase-1 activation, macrophage death, and extracellular bacterial release. A) Dot plots of viability staining (Live and Dead) and active caspase-1 staining in myeloid (Live/Dead⁻, DUMP⁻, CD45⁺) cells from the lungs of Mtb-infected WT and *Entpd1*^−/-^ mice. B-D) Percentages of live cells, live active caspase 1^+^ cells, and dead active caspase 1^+^ cells in mice described in A. E) Tunnel staining in WT and Entpd1-/-mice lung histological sections at day 21 of infection. F) Number of Tunnel^+^ cells/field in mice described in E. G) Mean Fluorescence Intensity (MFI) of Tunnel staining/field in mice described in A. Each dot represents the mean of six fields per mouse. H) Histograms of viability staining (Live and Dead) in bone marrow-derived macrophages (BMDM) from WT or *Entpd1^−/-^* mice after 24h post-infection *in vitro* with an MOI of 10. I and J) Percentage of dead cells and absolute number of live cells per well in BMDM cultures described in H. K) Extracellular CFU and percentage of LHD release in the supernatant of infected BMDM cultures described in H. Data represents two independent experiments. Significant differences between groups are indicated by p < 0.05. Significance levels are described as follows: p < 0.05 (*), p < 0.01 (**), p < 0.001 (***), and p < 0.0001 (****), determined by the Mann-Whitney non-parametric test.

To determine whether the caspase 1-associated myeloid cell death in infected *Entpd1^−/-^* lungs was linked to DNA fragmentation, we stained lung histological sections using the TUNEL method. Considering both the number of positive cells and the mean fluorescence intensity (MFI), a higher level of DNA damage was observed in infected lungs of *Entpd1^−/-^* mice compared to WT mice (Fig. 5E-G). This finding, combined with the increase of dead myeloid cells expressing high levels of active caspase-1+ (Fig. 5A), suggests an inflammasome-driven cell death, possibly via pyroptosis or secondary apoptosis, contributing to enhance DNA fragmentation when TB develops in the absence of CD39.

To specifically assess the role of CD39 in Mtb-induced macrophage killing, we compared bone marrow-derived WT and *Entpd1^−/-^* macrophages (BMDMs). The absence of CD39 resulted in 25% fewer viable BMDMs following infection (Fig. 5H and 5I) and a significant reduction in the absolute number of these cells (Fig. 5J). Suggesting that infected *Entpd1^−/-^* BMDMs were more susceptible to cell lysis, culture supernatants of these cells showed an increase in the numbers of extracellular mycobacteria and higher levels of lactate dehydrogenase (LDH), a marker of membrane integrity loss (Fig. 5K and 5I). These findings highlight the critical role of CD39 in regulating macrophage survival during Mtb infection and thus in limiting extracellular bacterial spread.

### P2RX7 inhibition attenuates the effects of CD39 deficiency *in vivo* and *in vitro*, promoting macrophage survival and preventing necrotic lesions and bacterial dissemination

It is known that CD39 can regulate macrophage death and the release of pro-inflammatory cytokines induced by P2RX7 signaling (19, 30). Therefore, we next investigated whether the protective effect of CD39 on Mtb-infected killing resulted from its ability to degrade eATP and prevent P2RX7 signaling. The inhibition of P2RX7 with A740003 was able to rescue approximately 35% of *Entpd1*^−/-^ BMDM viability after M299 infection using a MOI of 10 (Fig. 6A). A similar difference was observed in the absolute number of live cells per culture (Fig. 6B). As evidence of the protective effect of P2RX7 inhibition on necrotic death of these cells, a lower number of extracellular CFUs was determined in the supernatants of A740003-treated infected *Entpd1*^−/-^ BMDMs (Fig. 6C). These findings indicate that CD39 impairs P2RX7-triggered macrophage death induced by highly virulent mycobacteria.

**Figure 6.**
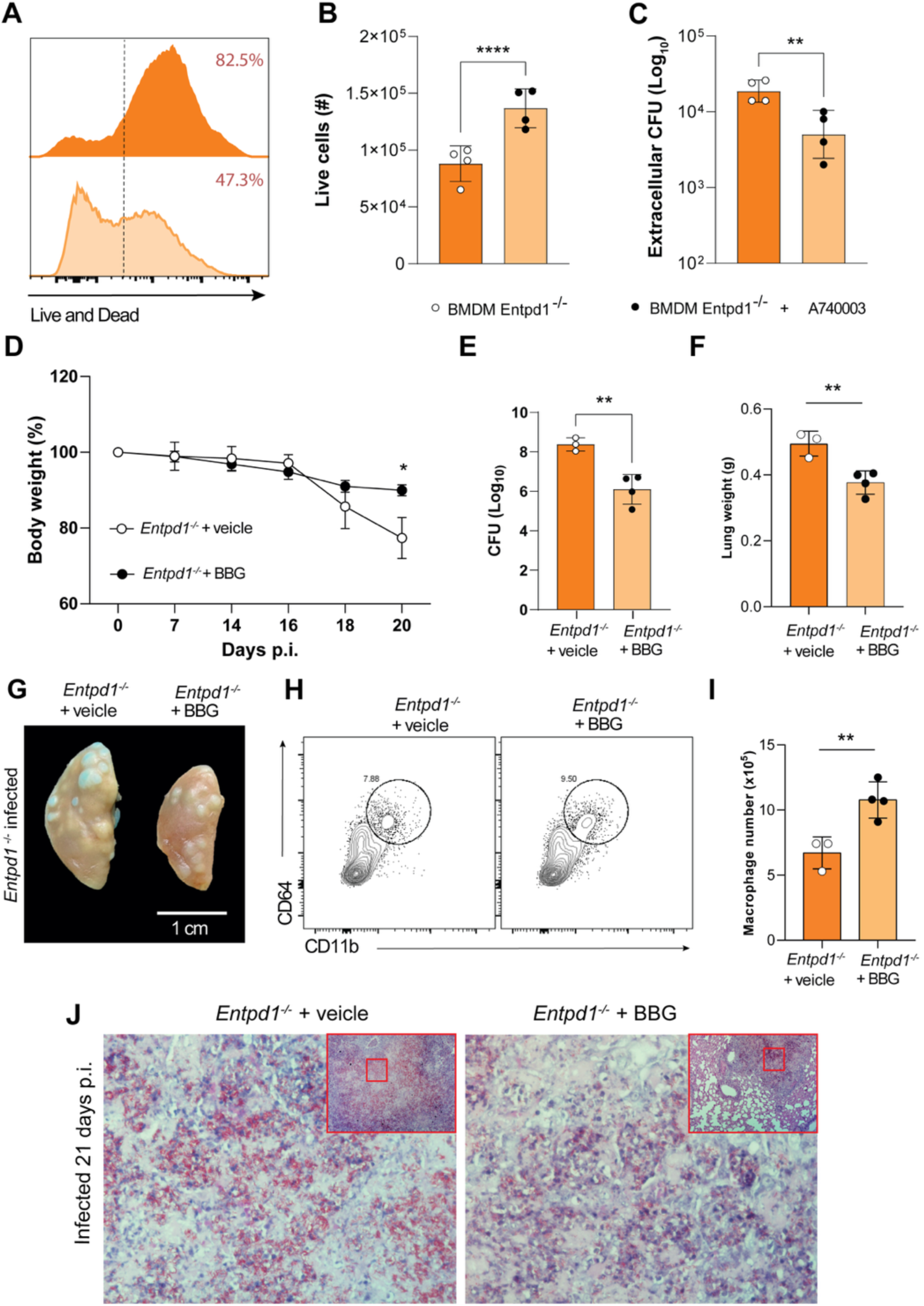
P2RX7 inhibition attenuates the effects of CD39 deficiency *in vivo* and *in vitro*. (A) Histograms showing viability staining (Live and Dead) of bone marrow-derived macrophages (BMDMs) from *Entpd1*^−/-^ mice treated or not with the P2RX7 inhibitor Brilliant blue G (BBG). Cells were maintained in culture for 24h post-infection (p.i.) with an MOI of 10. B) Absolute numbers of BMDMs in cultures described in A. C) Extracellular colony-forming units (CFUs) in the supernatants of infected BMDM cultures described in A. The data represents two independent experiments. D) Percentage changes in body weights of infected *Entpd1^−/-^* mice treated or not with BBG relative to baseline (day 0). E and F) CFUs of M299 Mtb in lung homogenates and lung weights of mice at 21 days of infection. G) Gross macroscopic pathology of the left lung of mice at 21 days of infection. H and I) Contour plots and absolute numbers of lung macrophages of mice at 21 days of infection. J) Histopathological analysis with Ziehl-Neelsen staining of lung sections of mice at 21 days of infection. Significant differences between groups are indicated by p < 0.05. Significance levels are represented as follows: p < 0.05 (*), p < 0.01 (**), p < 0.001 (***), and p < 0.0001 (****), determined by the Mann-Whitney non-parametric test. Data represents two independent experiments with three to four mice per group.

The critical role of CD39 in protecting against the development of necrotic lesions and bacterial spread in severe TB may also result from its ability to reduce the eATP sensing by P2RX7. To test this hypothesis, we inhibited P2RX7-signaling pharmacologically *in vivo*. *Entpd1*^−/-^ mice were treated with the P2RX7 inhibitor Brilliant Blue G (BBG) every two days from day 14 to 21 p.i. when the mice started to lose weight, as previously reported (14). At day 21 p.i., mice treated with the P2RX7 inhibitor showed less body weight loss compared to untreated animals (Fig. 6D). Importantly, inhibition of P2RX7 in *Entpd1*^−/-^ mice increased resistance to Mtb infection, as evidenced by reduced bacterial burden, lung weight, and the number and size of white nodules in the lung (Fig. 6E-G). Following these findings, the population of lung macrophages was increased after BBG treatment (Fig. 6H-I). In addition, because of P2RX7 inhibition, the histopathological analysis showed a significant reduction in pulmonary necrosis areas associated with extracellular bacteria (Fig. 6J). These results show that the beneficial effects of CD39 in suppressing the development of severe forms of TB in mice are primarily mediated by its ability to prevent eATP sensing by P2RX7.

## DISCUSSION

Necrotic pneumonia is a severe condition associated with pulmonary TB that has a significant impact on disease progression and transmission by promoting pathogen dissemination through lung tissue and respiratory tracts (31, 32). Despite advances in understanding the role of P2RX7 in the development of severe TB pathology (12, 14, 29, 33), important questions related to the regulation of this pathway remain to be answered. The ability of CD39 to degrade eATP and generate adenosine in the extracellular environment makes this molecule a possible target for regulating P2RX7 activation and the inflammatory response during TB (34, 35). Addressing this issue, our study reveals a major role for CD39 in TB progression by controlling P2RX7-induced death of infected macrophages and, consequently, reducing necrotic lesions in the lungs and pathogen dissemination.

The first indication of the involvement of CD39 in the regulation of P2RX7 signaling during TB was the elevated expression of *ENTPD1* and *P2RX7* genes, but not of genes associated with other ecto-ATPases and adenosine signaling (*ENTPD2,3*, *NT5E*, and *ADORA1,2B,3*) in peripheral blood cells of TB patients. Coexpression of genes encoding CD39 and P2RX7 was also observed in NHP granuloma cells, as well as in the blood and lung cells from Mtb-infected C57BL/6 mice. Since the expression *ENTPD1* and *P2RX7* genes was not upregulated in peripheral blood cells during latent human infection, these molecules represent potential biomarkers for identification, appropriate treatment and management of active TB patients. Stimuli associated with tissue damage, hypoxia (via hypoxia-inducible factor 1 (HIF1) transcription factor), and inflammation can upregulate CD39 and P2RX7 expression (15, 16, 36–40). Among them, cytokines with systemic activity, such as IL-1β, IL-6 and TNF, as well as type I IFNs, are produced during TB (41) and may be responsible for the increased expression of CD39 and P2RX7 in peripheral blood cells during active disease.

Excessive inflammatory responses were observed in the lungs of CD39-deficient mice, as evidenced by high levels of IL-1β in the lungs and robust activation of caspase-1 in myeloid cells. Of note, an increased population of dead myeloid cells exhibiting intense caspase-1 activity was found in the lungs of CD39-deficient mice, suggesting that CD39 protects these cells to undergoing necrosis associated with inflammasome activation, as it has been reported during pyroptosis (42). Supporting the concept that CD39 protects against tissue damage during TB, increased DNA fragmentation was observed in the lungs when the disease developed without CD39. Compelling evidence suggesting that CD39 limits TB progression by regulating P2RX7 activation comes from the comparative analysis of WT and CD39-deficient mice infected with hypervirulent mycobacteria and treated with a P2RX7 inhibitor. CD39 deficiency promoted widespread lung pulmonary necrosis and enhanced bacterial growth and dissemination, culminating in early loss of body weight and, consequently, increased mortality in mice. Significantly, pharmacological P2RX7 blockade attenuated the progression of the disease caused by the absence of CD39. A parallel finding has been described in sepsis in which P2RX7 also plays a significant role in disease development (19), as reported in severe TB (12). Interestingly, a similar phenotype was observed in an animal model of sepsis, in which CD39 deficiency promoted P2RX7-triggered cell death, cytokine production, and liver injury (19) .

Our results also showed that macrophages are critical players in the regulatory activity of CD39 in P2RX7-mediated development of necrotic lesions in pulmonary TB. Macrophages and endothelial cells of NHP granuloma exhibited high transcriptional levels of genes encoding CD39 and P2RX7. Still, these genes were expressed mainly in the macrophage population, considering their prevalence in this tissue. Concordantly, in the lungs of Mtb-infected C57BL/6 mice, CD39 and P2RX7 proteins were expressed at higher levels in immune cells than structure cells, particularly in macrophages that showed the highest expression of both molecules within immune cells. Remarkably, the lack of CD39 restricted to the immune cell compartment was sufficient to promote extensive necrotic lesions in the lungs, together with increased bacterial growth and dissemination in mice infected with hypervirulent mycobacteria. Importantly, a protective effect against severe TB was observed in mice repopulated with immune cells that did not express P2RX7 (33). It has been reported that macrophages infected with avirulent mycobacteria die through apoptosis, preventing bacterial dissemination, whereas virulent mycobacteria trigger necrosis of infected macrophage, thereby facilitating pathogen release into the extracellular environment (11, 43, 44). In this context, we have previously shown that P2RX7 signaling is critical in triggering necrotic death of macrophages infected with highly virulent mycobacteria (28). Extending these findings, this study demonstrated that CD39 regulates the necrotic death of infected macrophages by inhibiting P2RX7 signaling. CD39 deficiency in Mtb-infected mice resulted in a profound depletion of lung macrophages associated with increased numbers of dead cells and release of IL-1α and IL-1β in the lungs. Importantly, blocking P2RX7 in these animals increased the macrophage population in the lungs of these mice. Our *in vitro* results also showed that infected macrophages lacking CD39 were predisposed to necrotic death in a P2RX7-dependent manner. Since CD39 is known to regulate the balance between cell survival and death, we postulate that its deficiency leads to uncontrolled P2RX7 activation by eATP. This excessive activation promotes increased necrotic death of macrophages, compromising the host’s ability to control the infection. Furthermore, the release of intracellular contents amplifies inflammation and exacerbates tissue damage, creating a permissive environment for pathogen persistence and immune dysregulation.

Our findings strongly suggest that CD39 shapes the development of pulmonary TB by acting as a counterregulatory mechanism that limits P2RX7 signaling in infected macrophages via eATP hydrolysis. Thus, CD39 protects these cells from necrotic death, maintaining their immune response function and preventing the disease progression. Altogether, these results provide further evidence for the critical role of purinergic signaling in regulating cellular/tissue damage as well as disease progression and implicate the CD39/P2RX7 axis as a potential target for host-directed TB therapy.

## MATERIALS AND METHODS

### Mice

Specific pathogen-free C57BL/6 (WT) and *Entpd1^−/-^* male (6-8-week-old) mice were bred at the isogenic mouse facility, ICB, USP. After infection, mice were maintained in micro isolator cages with HEPA filters at the Biosafety Laboratory Level 3, FCF, USP. All procedures were performed according to national regulations and ethical guidelines for mouse experimentation (permit no. 3185080318).

### Mycobacteria and mouse infection

The Mtb strain of the Beijing genotype (strain M299), isolated from a TB patient in the Maputo province, Mozambique, was kindly provided by Dr. Philip Suffys (Oswaldo Cruz Foundation, FIOCRUZ, Rio de Janeiro, Brazil). Frozen aliquots were thawed and grown in Middlebrook 7H9 medium enriched with 10% (vol/vol) ADC (albumin, dextrose, catalase) (Difco, BD Biosciences, USA) and 0.05% (vol/vol) Tween 80 (Sigma-Aldrich) and maintained at 37°C for seven days until mid-log phase in constant agitation. The bacterial suspensions were sonicated in a water bath and vortexed for 1 minute to disperse the lumps. The density of bacterial suspensions was determined using a spectrophotometer at 600 nm. Mice were anesthetized intraperitoneally (i.p.) with ketamine (Vetbrands, Brazil; 100 mg/kg) and xylazine (Vetbrands; 15 mg/kg) and infected intratracheally (i.t.) with ∼70-100 (lower dose) or ∼150-200 (double dose) bacilli of the highly virulent Mtb Beijing M299 strain (28).

### P2RX7 inhibition to prevent cell death during lung processing

Several studies describe that P2RX7 direct, or indirect activity inhibition preserves cell viability during tissue processing under collagenase IV digestion (45, 46). To prevent cell death during lung digestion and processing to obtain cell suspensions for Flow Cytometry, infected mice were injected intravenously with brilliant blue G (BBG, Sigma-Aldrich), a P2X receptor inhibitor (45 mg/Kg/mouse in 200 µL of PBS) 30 minutes before the euthanasia (**Supplementary Figure 1A**). The differences in the cell viability between treated and non-treated infected mice prior to euthanasia are shown in **Supplementary Figures 1B-G**.

### Isolation and counting of lung-infiltrating cells

The dissected lung lobes were washed with sterile PBS 1x, fragmented, and digested with collagenase type IV (0.5 mg/mL, Sigma-Aldrich) and type IV bovine pancreatic DNAse (Roche Diagnostics; 1 mg/ml) in RPMI 1640 medium (Gibco, USA) at 37°C for 40 minutes under agitation (200 rpm) (Almeida *et al.*, 2017). The remaining lung fragments were mechanically dissociated by passage through a 100 µm pore-size cell strainer and incubated with ACK Lysing Buffer (Thermo Fisher Scientific, USA) at room temperature for one minute to deplete the erythrocytes. Cell suspensions were washed with 10% fetal calf serum (FCS, Gibco) in PBS following centrifugation at 1,200 rpm for 5 minutes and resuspended in RPMI 1640 medium. Lung cell viability was determined using a trypan blue exclusion assay and a hemocytometer.

### Phenotypic analysis of lung-infiltrating cells

Isolated lung cells (1×10^6^ cells/well) were seeded in round-bottom 96-well plates and stained with Live/dead dye (Thermo Fisher Scientific) to determine cell viability, as described in the datasheet. Lung cells were stained using fluorochrome-labeled monoclonal antibodies to lineage (CD4 -RM4-5, CD8 - S3-6.7, CD19 - 1D3 and NK.1 - PK136), CD11b (M1/70), Ly6G (1A8), Ly6C (AL-21), CD11c (N418), F4/80 (BM-8), CD39 (24DMS1), CD4 (RM4.5), CD8 (S3-6.7), CD44 (IM7) and CD69 (H1.2F3) (BD Biosciences) for 30 minutes at 4°C. Cells were fixed with 4% paraformaldehyde for 30 minutes at 4°C and washed on a staining buffer. Cell acquisition was performed using the LSRFortessa™ flow cytometer (BD Biosciences – USA) and the FlowJo 10.5.3 software (BD Biosciences). The gating strategy for CD11b^+^ myeloid cell analysis is shown in Supplementary Figure 1H.

### Lung macroscopic and microscopic analyses

The harvested lung lobes were washed with sterile PBS 1x and weighed. The lung relative mass was calculated by dividing the mean lung weight in experimental mice by the mean in uninfected controls from their respective groups. The left lung upper lobe was maintained in 10% buffer formalin for 48 hours, photographed, and subsequently embedded in paraffin. Histological sections of approximately 4-5 μm were stained using the hematoxylin-eosin (HE) method to visualize tissue alterations, Ziehl Neelsen (ZN) method to detect the presence of acid-fast bacteria (AFB), and Immunofluorescence for TUNEL Staining (Thermo Fisher Scientific), as described in the datasheet, to visualize nucleus fragmentation. The samples were examined with an Axioplan microscope (Carl Zeiss Inc., Germany), and the images of lung sections were captured by Coolpix P995 (Nikon)-coupled device camera.

### Colony-forming unit (CFU) analysis and counting

Corresponding bacterial concentrations in lung homogenate were determined by the colony-forming unit test, using serial 10-fold dilutions of each suspension and plating on Middlebrook 7H10 agar (Difco, Detroit, MI), supplemented with 0.5% glycerol, 10% oleic acid–albumin-dextrose–catalase enrichment, OADC (BD, Sparks, MD). Plates were cultured at 37°C for 21 days, and total CFU was determined by colony counting.

### BMDM cultures

Murine BMDMs were generated by isolating undifferentiated monocytes from bone marrow from both femurs and tibiae of WT and *Entpd1^−/-^* mice. Bone marrow was harvested in DMEM/F-12 (Gibco) supplemented with 10% heat-inactivated FBS (Gibco) and flushed through a syringe with a 16-gauge needle. Cells were dispersed with a 5-mL syringe (BD Biosciences) fitted with a 20-gauge needle. Dispersed cells were seeded in culture flasks (T-175) containing 20 mL DMEM/F-12 supplemented with 10% FBS, 25 µg/ml gentamicin (Gibco), and 20% of L929-conditioned media. Cells were incubated at 37°C with 5% CO2, and 20 mL of fresh medium containing L929-conditioned media without gentamicin was added on day 3. On day 7, macrophages were detached by the addition of cold PBS.

### *In vitro* BMDM infection

Frozen bacterial aliquots were thawed and cultured as described above. Bacterial suspensions were centrifuged at 4,000 rpm for 10 min, resuspended in DMEM/F-12, sonicated for 30 s, and homogenized to reduce bacterial clumping. BMDMs were exposed to M299 Mtb infection at 1:10 MOI. After 3 hours, cells were washed three times with room temperature PBS and then cultured in fresh DMEM/F-12 media, supplemented with 3% for 24 hours. According to the manufacturer’s instructions, LDH release in the supernatants from BMDM cultures was determined using the CytoTox 96 nonradioactive cytotoxicity assay (Promega).

### Statistical analyses

Statistical analyses were performed using the GraphPad Prism 7 software (GraphPad, USA), and differences between groups were considered significant when p < 0.05 (5%). ANOVA and Bonferroni post-test analyzed the simultaneous effects of two factors. One-way ANOVA and Tukey test were used to evaluate the impact of a single parameter.

## NOTES

## Acknowledgements

We thank Dr. Philip Suffys (Oswaldo Cruz Foundation, FIOCRUZ, Rio de Janeiro, Brazil) for providing the Beijing M299 *M. tuberculosis* strain. We also thank Rogério Silva do Nascimento, José Israel Lima, Silvana Silva and Maria Áurea de Alvarenga for technical assistance.

## Financial support

This study was supported in whole by São Paulo Research Foundation (FAPESP-Brazil) grants: 2015/20432-8 (M.R.D.L.), 2019/24700-8 (I.S.C.), and 2020/09043-8 (C.C.B.B.); and by National Council for Scientific and Technological Development (CNPq) grants: 408909/2018-8 (M.R.D.L.), 303810/2018-1 (M.R.D.L.), 308870/2023-0 (M.R.D.L.) and 140666/2018-4 (G.A.S.).

## Authors’ Contributions

Conceived and designed the experiments: G.A.S., E.P.A. E.L. and M.R.D.L. Performed the experiments: G.A.S.; I.S.C.; F.M.A.; C.C.B.B.; D.G.C; C.R.S.; B.G.M.; P.H.L.R.; J.T.X.J; and Analyzed the data: G.A.S.; F.M.A.; J.C.S.S; E.L. and M.R.D.L. Contributed reagents/materials/ analysis tools: G.A.S.; M.S.R; J.M.A.; M.H.H.; R.C.S.; S.C.R.; E.L.; M.R.D.L.

## Potential conflicts of interest

The authors have declared no conflicts of interest.

This study has not been previously reported and is not being considered for publication elsewhere. All authors agree to submit this manuscript.

## SUPLEMENTARY MATERIAL

**Supplementary Figure 1.**
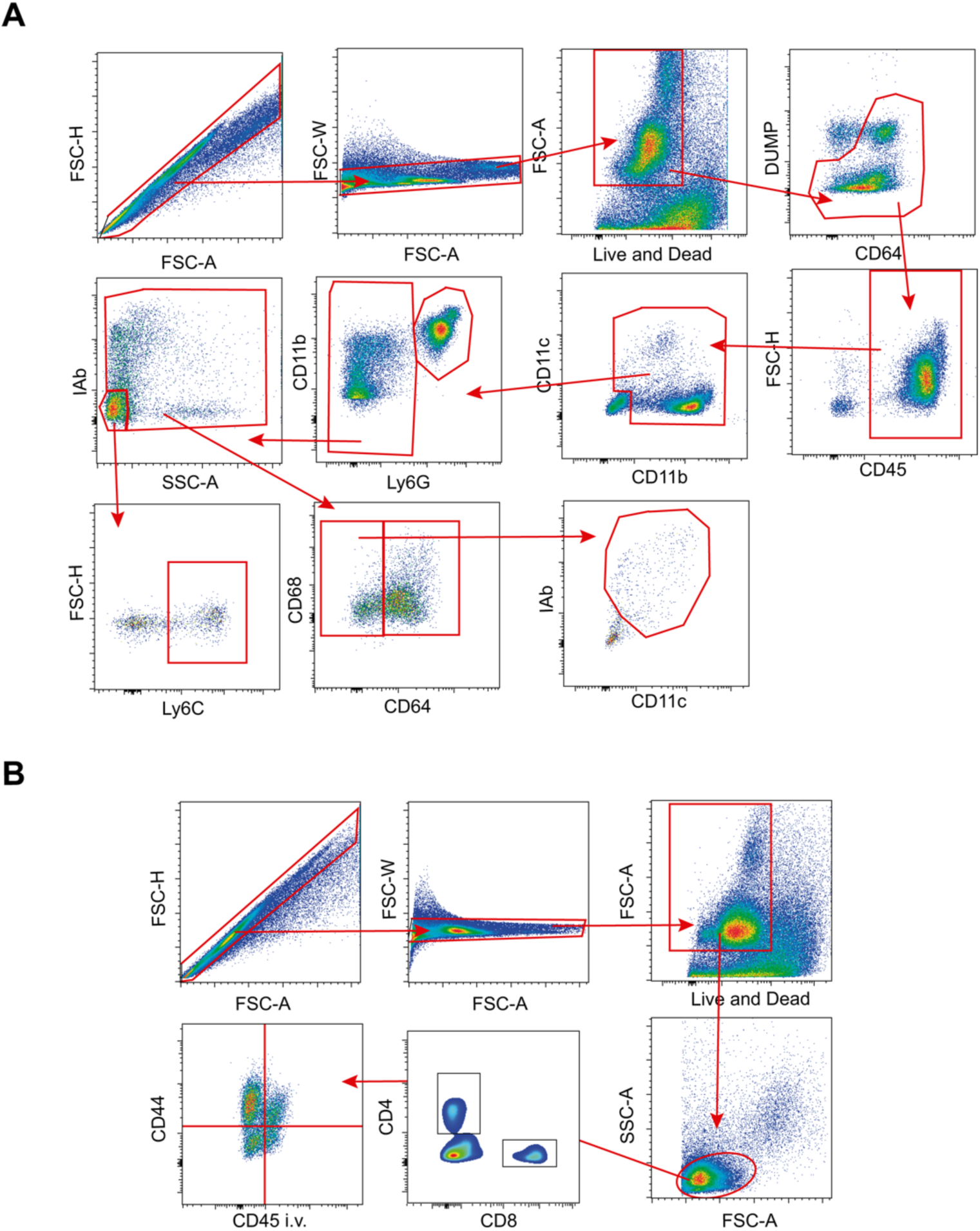
Gate strategies for analyzing myeloid cells and CD4+ T cells are shown in mice infected with the M299 Mtb strain. (A) Gate strategies to evaluate macrophages (Live/Dead⁻, DUMP⁻, CD45⁺, Ly6G⁻, CD68⁺, CD64⁺), dendritic cells (Live/Dead⁻, DUMP⁻, CD45⁺, Ly6G⁻, CD68⁻, CD64⁻, CD11b⁺, MHCII⁺), monocytes (Live/Dead⁻, DUMP⁻, CD45⁺, Ly6G⁻, MHCII⁻, SSCLow, Ly6C⁺), and neutrophils (Live/Dead⁻, DUMP⁻, CD45⁺, Ly6G⁺) within the CD45⁺ DUMP^−^ population are shown. (B) Gate strategies are shown to evaluate CD8^+^, CD4^+^ T cells, and i.v.^+^ (vascular) and i.v.^−^ (parenchymal) CD44^+^CD4^+^ T cells.

**Supplementary Figure 1.**
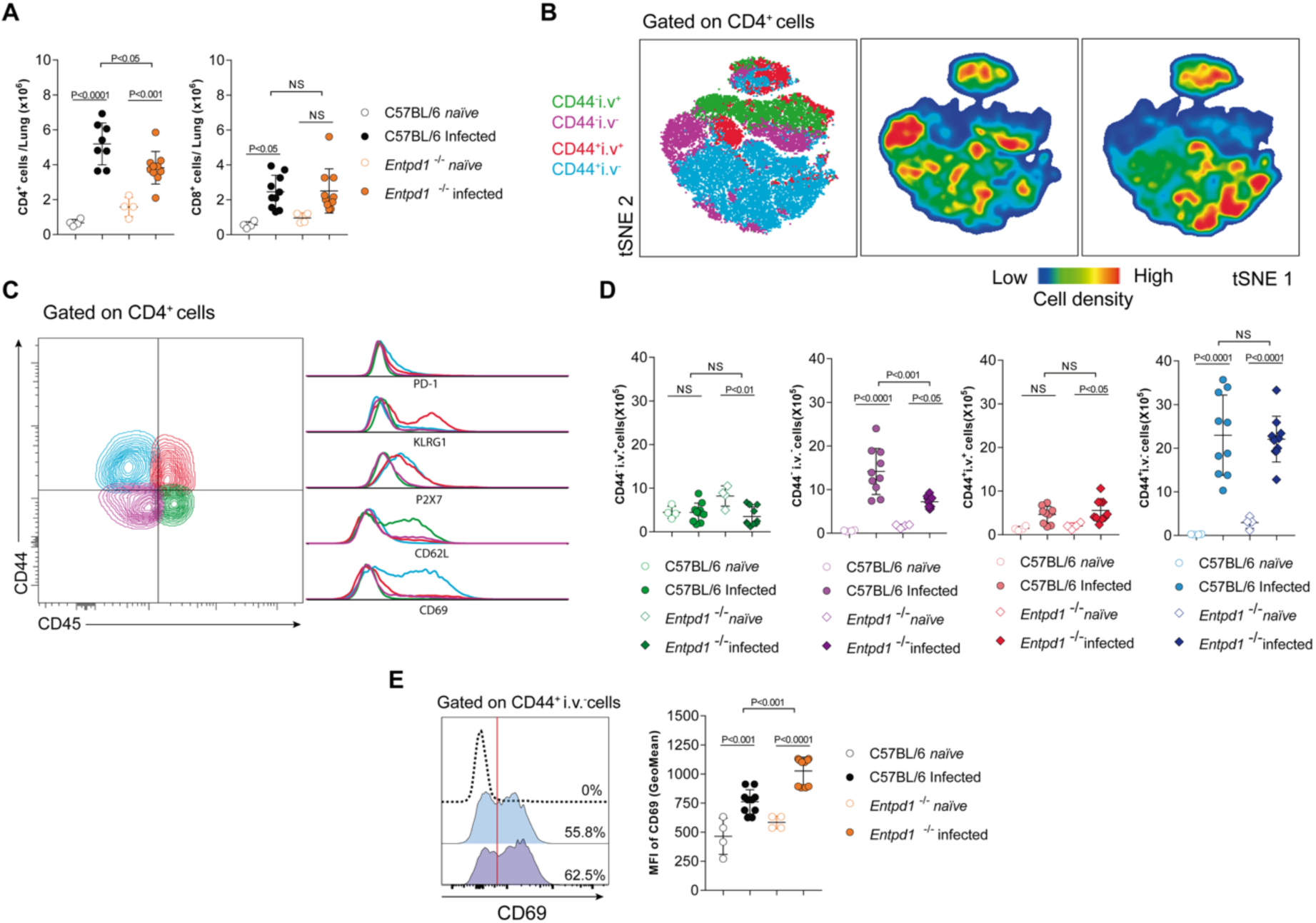
CD39 deficiency does not affect the number of activated CD4 T cells in the lung parenchyma or vasculature. A) Numbers of CD4^+^ T cells. B) Numbers of CD8^+^ T cells. C) tSNE plots depict the distribution of vascular and parenchymal CD4^+^ T cell subsets. C) Expression of markers that are expressed by those distinguished subsets. D) Numbers of i.v.^+^ (vascular) and i.v.^−^ (parenchymal) CD44^+^CD4^+^ T cells and CD44^−^CD4^+^ T cells.

**Supplementary Figure 2.**
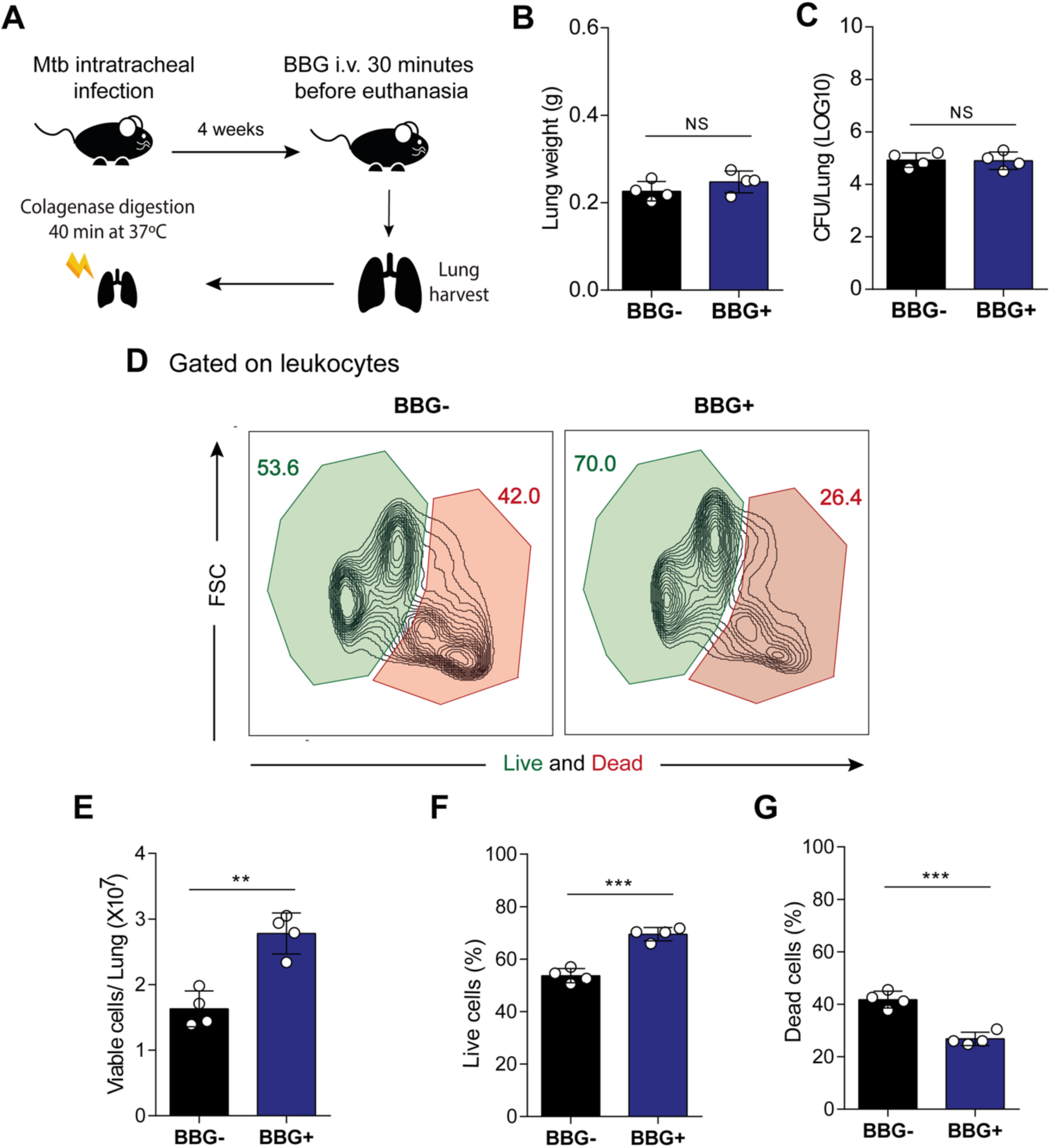
The strategy of inhibition of P2RX7 to prevent cell death during tissue processing to obtain cell suspensions. A) Experimental design: 4 weeks post-infection, Mtb-infected mice were treated or not with Brilliant Blue G (BBG) 30 minutes before euthanasia. After 30 minutes of blockade, the mice were euthanized, and the lungs were collected for digestion in a Collagenase + DNA cocktail. B) Lung weight and C) Colony-forming units (CFU) of Mtb M299 in lung homogenates from infected mice. D) Representative plots of live and dead cell frequencies on treated (BBG+) and non-treated (BBG-) mice. E) Number of viable cells per lung. F) Frequency of live cells. G) Frequency of Dead cells. Significant differences between groups are indicated by p < 0.05. Significance levels are represented as follows: p < 0.05 (*), p < 0.01 (**), p < 0.001 (***), and p < 0.0001 (****), determined by the Mann-Whitney non-parametric test. The data represents results from two independent experiments with four mice per group.

## Notes

### Competing Interest Statement

The authors have declared no competing interest.

